## Supplemental Figures and Legends for "Sexual dimorphism in melanocyte stem cell behavior reveals combinational therapeutic strategies for cutaneous repigmentation"

### Supplemental Figure 1

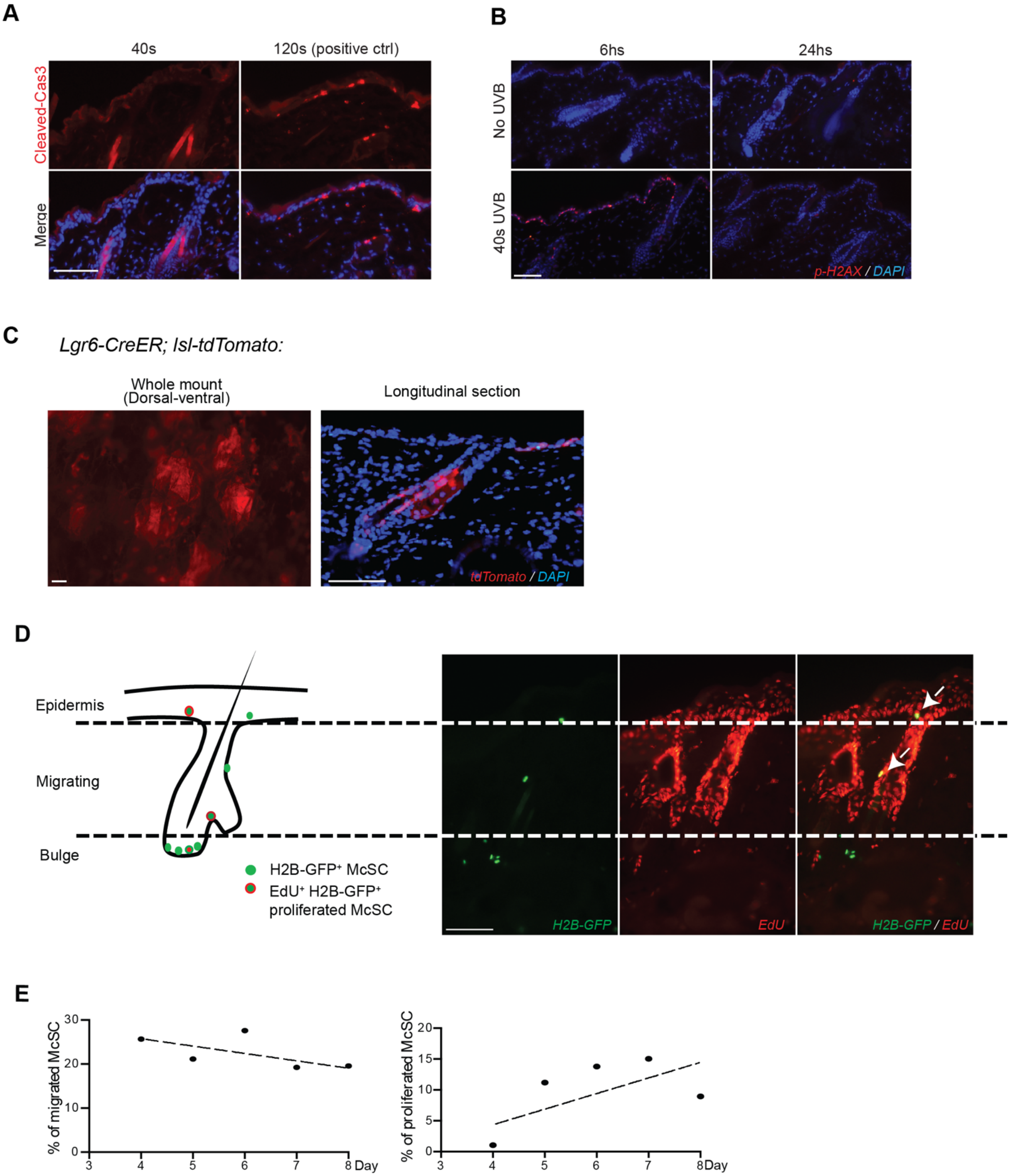

**Supplemental Figure 1**

(**A**) Representative images of cleaved-Caspase3 staining on control dorsal skin 24hs post UVB irradiation. n=3. Skin exposed to 120s of UVB irradiation was used as a positive control to show apoptosis. (**B**) Representative images of p-H2AX staining on UVB-irradiated dorsal skin at 6hs and 24hs. No-UVB skin was used as a negative control. As shown, DNA damaged cells were located on the surface (epidermis) at 6hs post first UVB irradiation, and most of the DNA damage had been resolved at 24hs post first UVB irradiation. n=3 for each time point. (**C**) Representative whole mount and longitudinal section images from *Lgr6-CreER; lsl-tdTomato* skin. As shown, tdTomato positive cells near the sebaceous gland do not come into focus via whole mount imaging. (**D**) Schematic of the bulge, migrating McSCs, and epidermal melanocytes (left). Representative images of McSCs labeled with EdU (right). Dashed lines denote the boundaries defining migrating melanocytes. $EdU^{+}H2B\text{-}GFP^{+}$ cells labeled by arrows indicate proliferated McSCs that were located in the middle of hair follicle (migrating) and located in epidermis (migrated). n>5. (**E**) Quantification of migrating melanocytes and melanocytes that incorporated EdU. Each dot represents one mouse. McSCs in 90-120 hair follicles were analyzed from each mouse.

### Supplemental Figure 2

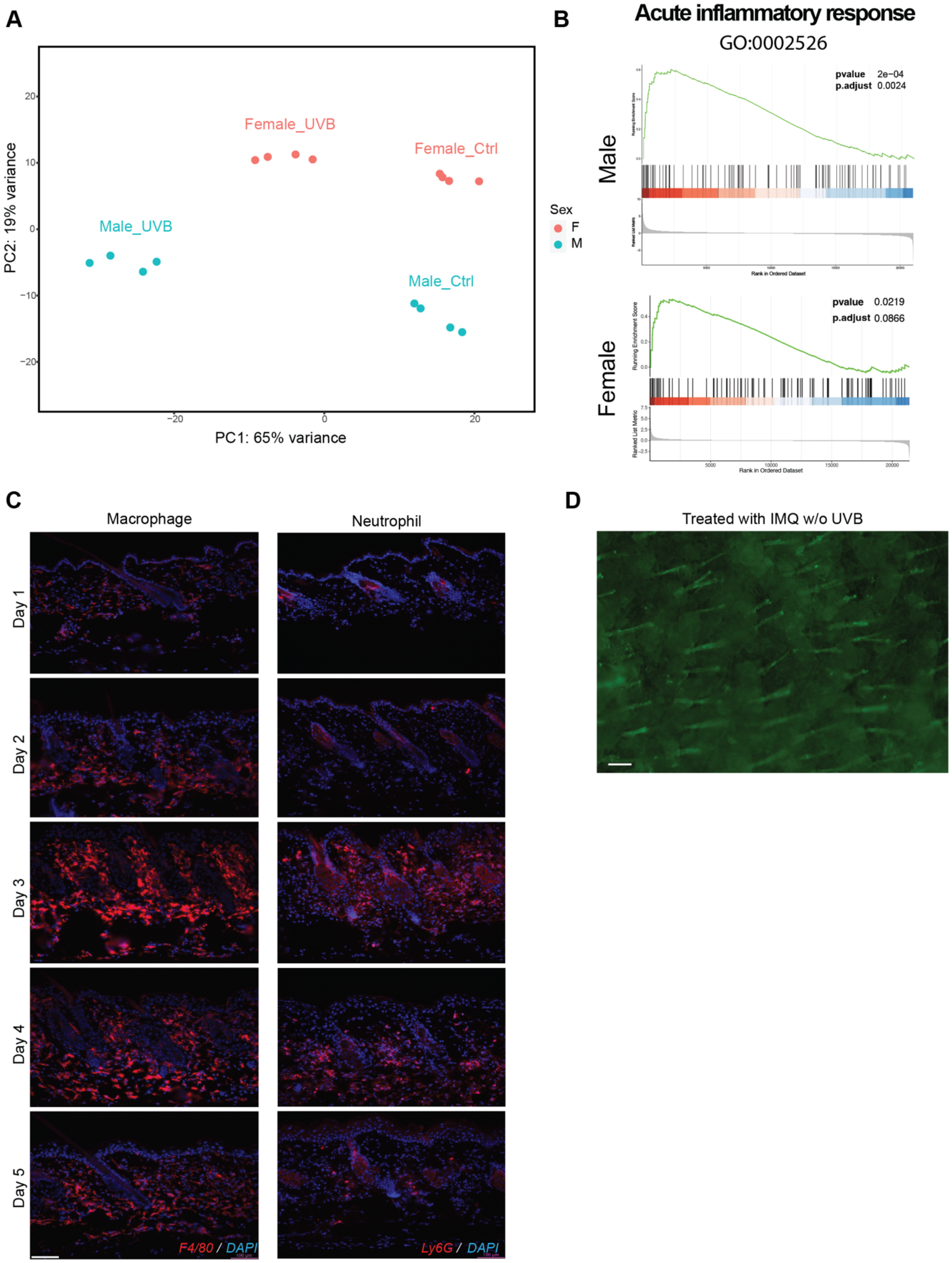

**Supplemental Figure 2**

(**A**) PCA plot of all 16 bulk mRNA-seq samples. Arrows indicate the transcription profile changes after UVB irradiation within each gender. Since a noticeable difference was found between genders, instead of directly comparing male UV samples with female UV samples, we compared the level of transcription change following UVB irradiation between genders. (**B**) Gene Set Enrichment Analysis (GSEA) of Acute Inflammatory Response (GO:0002526) on male and female samples. p.adjust value in female sample is 0.0866>0.05, not significant. (**C**) Representative images of F4/80 (Macrophages) and Ly6G (Neutrophils) staining on no-UVB skin and irradiated skin from day 2 to day 5. At least 3 male mice examined for each timepoint. (**D**) Representative images of IMQ treated skin without UVB irradiation. n=4 (2 males, 2 females). Scale bar: 100um.

### Supplemental Figure 3

**A**

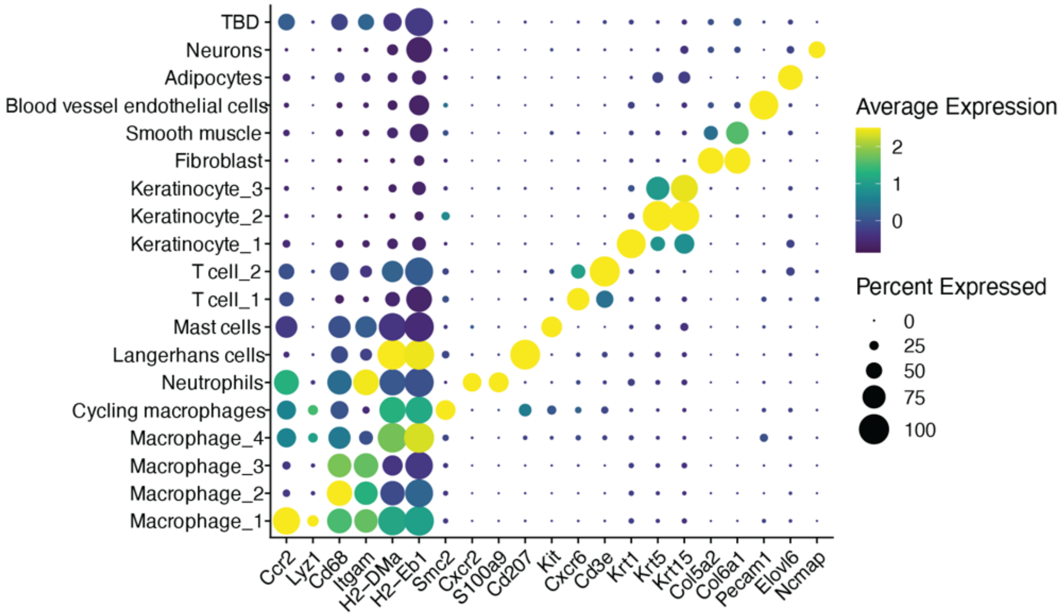


**B**

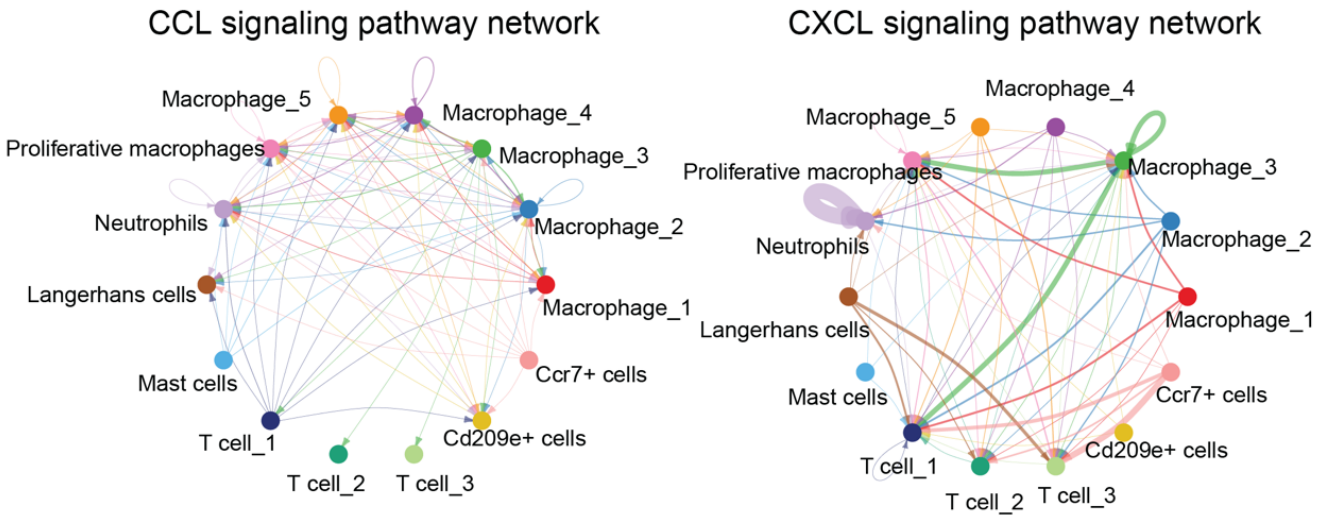

**Supplemental Figure 3**

(**A**) Dot plot showing the marker genes of each cell cluster. TBD (To Be Determined): unidentified cell cluster. (**B**) CellChat signaling analysis on CCL and CXCL signaling pathway on WT_UV sample. Arrows show the potential cell-cell communication and arrowheads show the signal receivers.

### Supplemental Figure 4

**A**

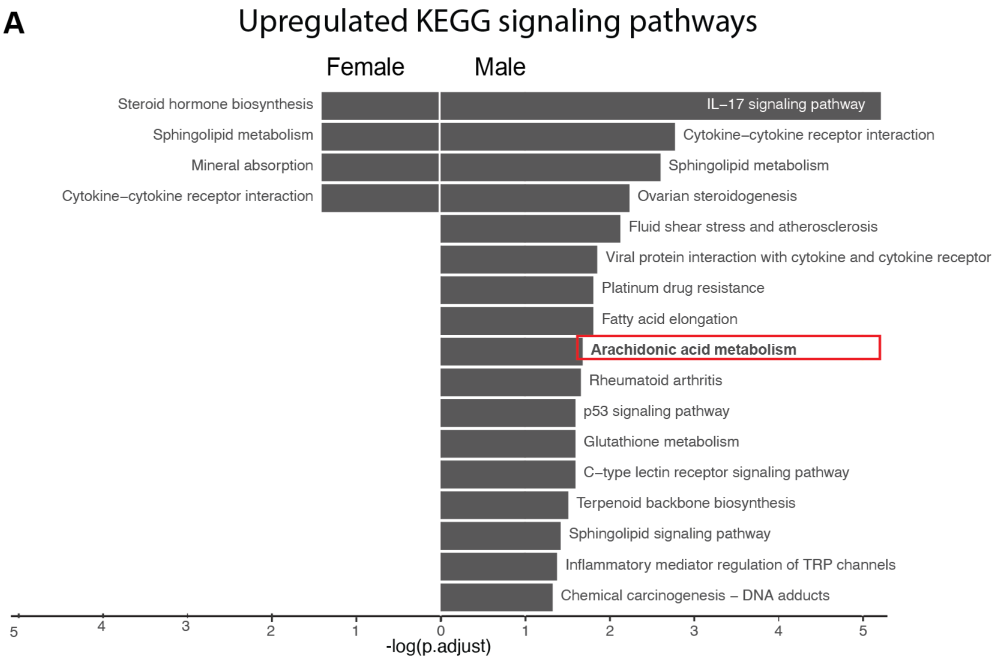

**Supplemental Figure 4**

(**A**) Upregulated KEGG signaling pathways in UVB-irradiated skin compared to no-UVB controls from female and male samples. Cutoff: pvalue $< 0.05$, qvalue $< 0.1$, padjust method is Benjamini & Hochberg.

### Supplemental Figure 5

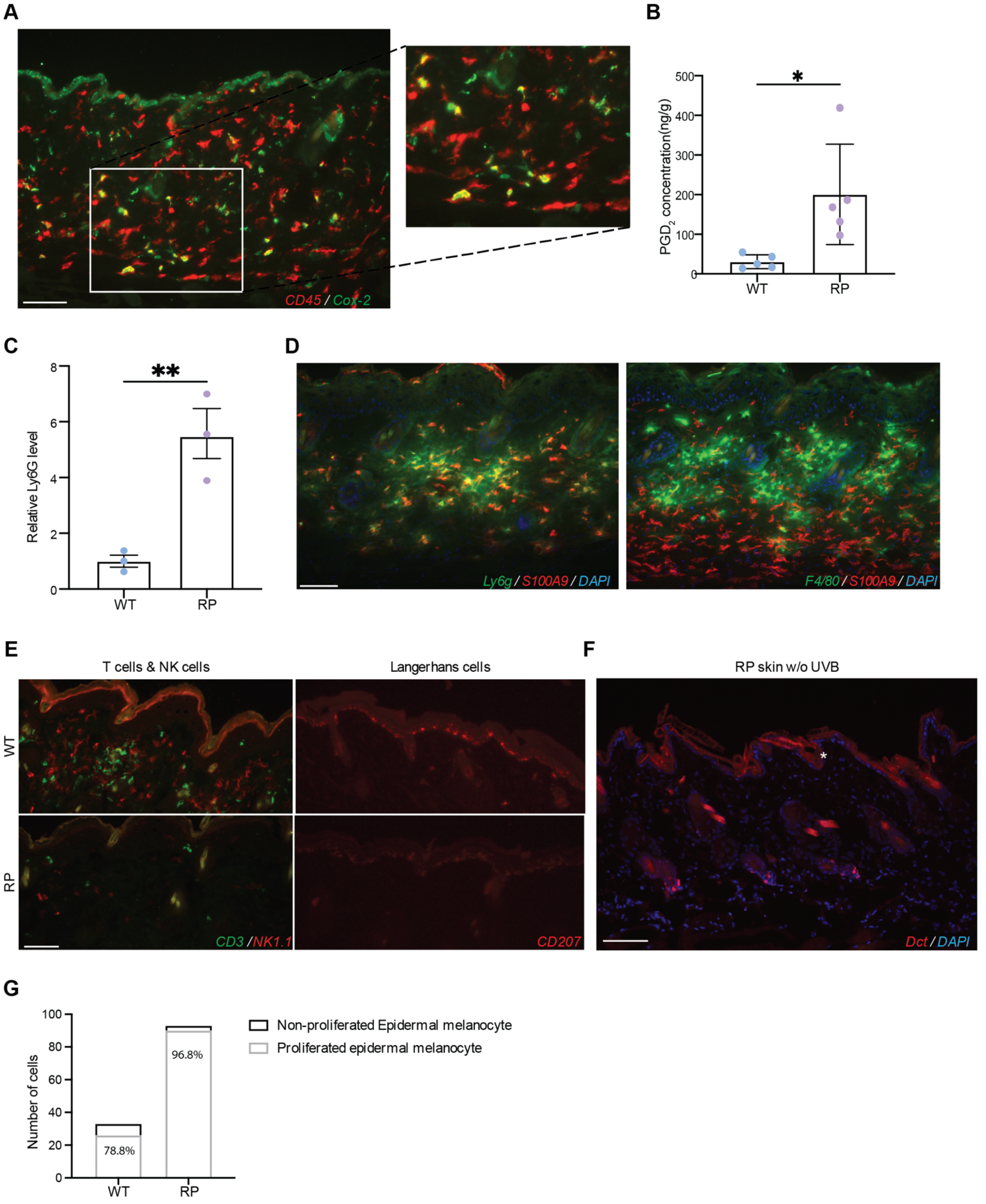

**Supplemental Figure 5**

(**A**) Representative images of CD45 and Cox-2 co-labeling in the dermis of RP mice. n>3. (**B**) ELISA quantification of skin tissue $PGD_2$ levels between Ctrl and RP mice collected at day 5. $PGD_2$ level was normalized by the tissue weight. Each dot indicates the normalized $PGD_2$ level in one animal. n=5 mice in each group. (**C**) Immunofluorescence staining quantification between Ctrl and RP mice collected at day 5. Relative Ly6G levels were quantified by the Ly6G positive staining area normalized by the DAPI area. At least 20 images were quantified in each mouse. (**D**) Representative images in RP mice collected at day 5 showing the co-localization between Ly6G neutrophils with S100A9 but not F4/80 macrophages, n=3 males. (**E**) Representative images of T cells (CD3), NK1.1 (NK cells), CD207 (Langerhans cells) in Ctrl and RP mice collected at day 5. As shown, significantly fewer T cells, NK cells and Langerhans cells were found in RP skin compared with Ctrls. n=3 males for each sample. (**F**) Representative images of no-UVB irradiated RP mice. No migrated melanocytes were found in the epidermis. Red staining in the image is background in the stratum corneum (labeled by *). n>5 males and 5 females. (**G**) Total number of migrated melanocytes in Ctrl and RP mice, and the percentage of proliferated melanocytes within the total migrated melanocytes. Control animals used in B,C were either single allele *Rosa-rtTA* or *Tre-Ptgs2* mice with doxycycline water treatment. Scale bar = 100um. Statistics: Welch's t-test, Error bar: SEM. *,** indicates pvalue<0.05 and 0.01 (**B,C**).

### Supplemental Figure 6

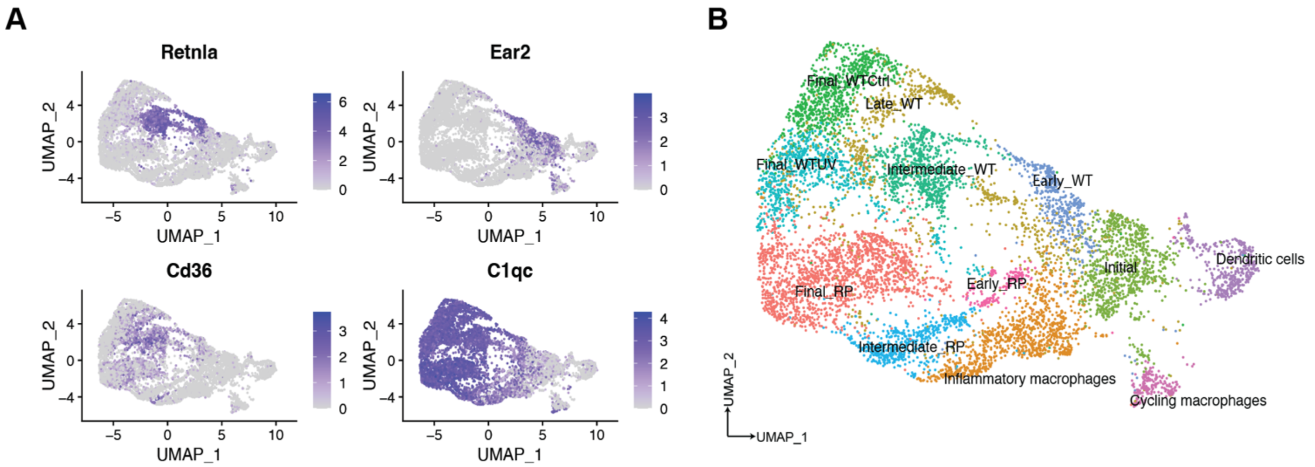

**Supplemental Figure 6**

(**A**) Visualization of early-stage macrophage marker *Retnla* and *Ear2*, and phagocytotic macrophage marker *Cd36* and *C1qc* on all macrophage sub-clusters from WT_Ctrl, WT_UV and RP_UV samples. (**B**) Sample group of macrophages with labeling based on the marker gene expression described in (**A**).

### Supplemental Figure 7

**A**

Treated with PBS w/o UVB

Treated with $dmPGE_2$ w/o UVB

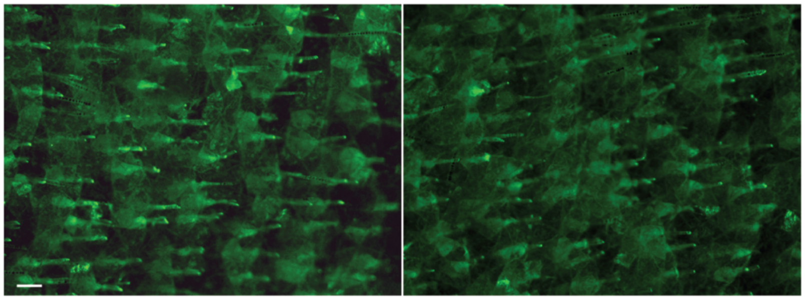

**B**

Treated with ruxolitinib w/o UVB

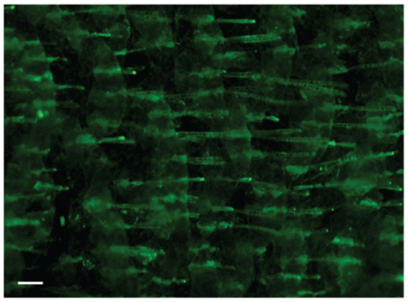

**Supplemental Figure 7**

(**A**) Representative images of no-UVB dorsal skin with and without $dmPGE_2$ treatment. No migrated melanocyte was found in $dmPGE_2$ treated mice in the absence of UVB. n=5 male mice. (**B**) Representative images of no-UVB dorsal skin with ruxolitinib treatment. n=4 female mice. Scale bar = 100um.
